# Protein interactome homeostasis through an N-recognin E3 ligase is a vulnerability in aneuploid cancer

**DOI:** 10.1101/2023.05.04.539299

**Authors:** Meena Kathiresan, Sambhavi Animesh, Robert Morris, Johannes Kreuzer, Krushna C. Patra, Lei Shi, Joshua Merritt, Xunqin Yin, Cyril H. Benes, Nabeel Bardeesy, Wilhelm Haas

## Abstract

Aneuploidy and resulting gene copy number alterations (CNAs) are important hallmarks of human cancers. Since CNAs are not associated with dosage compensation in mRNA expression, cancer cells with a high CNA burden must harbor mechanisms to mitigate proteotoxic stress resulting from stoichiometric imbalance and accumulation of unfolded proteins (*1*). Here, we show that aneuploid human cancer cells exhibit discordance between CNAs and protein levels due to compensation at the proteome level, mainly concerning multi-protein complexes. Moreover, we identify the N-recognin ubiquitin ligase UBR4 as a critical mediator of protein interactome homeostasis that is essential for viability, specifically in highly aneuploid cancers *in vitro* and *in vivo*. UBR4 prunes the proteome to ensure the balanced expression of protein complex members. Inactivation of UBR4 in highly aneuploid cancer cells causes a convergence of copy number and protein levels and induces proteotoxic stress pathways. UBR4 inhibition may present a broadly applicable therapeutic strategy for cancer and other diseases driven by aneuploidy.

**One-Sentence Summary:** The N-recognin ubiquitin ligase UBR4 as a critical mediator of protein interactome homeostasis that is essential for viability in aneuploid cancers.

---

Aneuploidy, defined as an aberrant number of whole chromosomes or chromosome arms, is the major cause of gene copy number alterations (CNA) in human cancer (*2, 3*). Increased aneuploidy is associated with changes in cancer pathophysiology—such as the transition from local to invasive disease (*4*) and from clonal to heterogeneous tumors (*5*)—and is a biomarker of worse prognosis (*6*). Whether aneuploidy represents a cause or a consequence of malignant transformation remains unclear. Regardless, the remarkable commonness of aneuploidy in cancer makes aneuploidy-related vulnerabilities a promising target for broadly applicable cancer therapies (*3*).

Cells with a high level of aneuploidy experience a range of cellular stresses, including proteotoxic stress that results from the stoichiometrically imbalanced expression of multi-protein complex members (*1*). This imbalance may lead to overstraining of the protein quality control machinery and the accumulation of toxic unfolded proteins as corroborated by the sensitization of aneuploid yeast strains to proteasome inhibition (*7*). Studies across a range of aneuploid systems, including yeast (*8*) and human tumors (*9*), have shown no widespread dosage compensation of CNA at the level of RNA expression. However, in disomic yeast strains, CNA dosage attenuation has been observed for ∼20 % of proteins – mainly members of multi-protein complexes (*10*). Furthermore, recent studies of human tumors and cancer models have shown a widespread discordance between CNA and protein levels (*9, 11, 12*) suggesting a highly efficient mechanism that prevents the toxic accumulation of stoichiometrically imbalanced and unfolded proteins in cancer. In a prior study of the proteome of > 40 breast cancer cell lines, we observed that co-regulation of protein concentrations across cell lines is a powerful predictor of protein-protein interactions, whereas RNA co-expression profiles exhibit little correlation with known interactions (*11*). This work demonstrated that large-scale profiling of protein co-regulation efficiently maps interactomes and their dynamics and has the potential to identify specific cancer vulnerabilities (*11*). In the present study, we aimed to understand the cellular mechanism that mediates the decoupling of CNA and protein levels, resulting in the accurate adjustment of concentrations of protein complex members in cancer.

Fig. 1a presents our working model for proteome homeostasis in aneuploid cancer cells: gene CNA are accurately reflected in the levels of transcribed RNA molecules. Upon translation, proteins quickly assemble into quaternary structures, while excess proteins need to be degraded to avoid proteotoxicity. The model predicts a high positive correlation between CNA and mRNA but not protein levels. Conversely, among constituents of multi-protein complexes, protein but not RNA levels should be co-regulated. Thus, the functional association between proteins correlates with suppression of the effect of CNA on their concentrations. We confirmed the model by examining our proteomics dataset representing 36 breast cancer cell lines (*11, 13*). For protein complexes listed in the CORUM database (*14*) we found a significantly higher correlation between CNA and mRNA than between CNA and protein levels (Fig. 1b, *P*<10^-15^, two-sample Kolmogorov-Smirnov test, table S1). Further corroborating the model, intra-protein complex correlation was significantly higher for protein than for mRNA levels (Fig. 1c, *P*<10^-15^, Kolmogorov-Smirnov, table S1). It should be noted that the data do not allow distinction between a protein synthesis or a protein degradation-based mechanism to adjust concentrations of protein complex members (*15*). However, this ambiguity does not affect our unbiased approach to investigating the mechanism.

**Figure 1.**
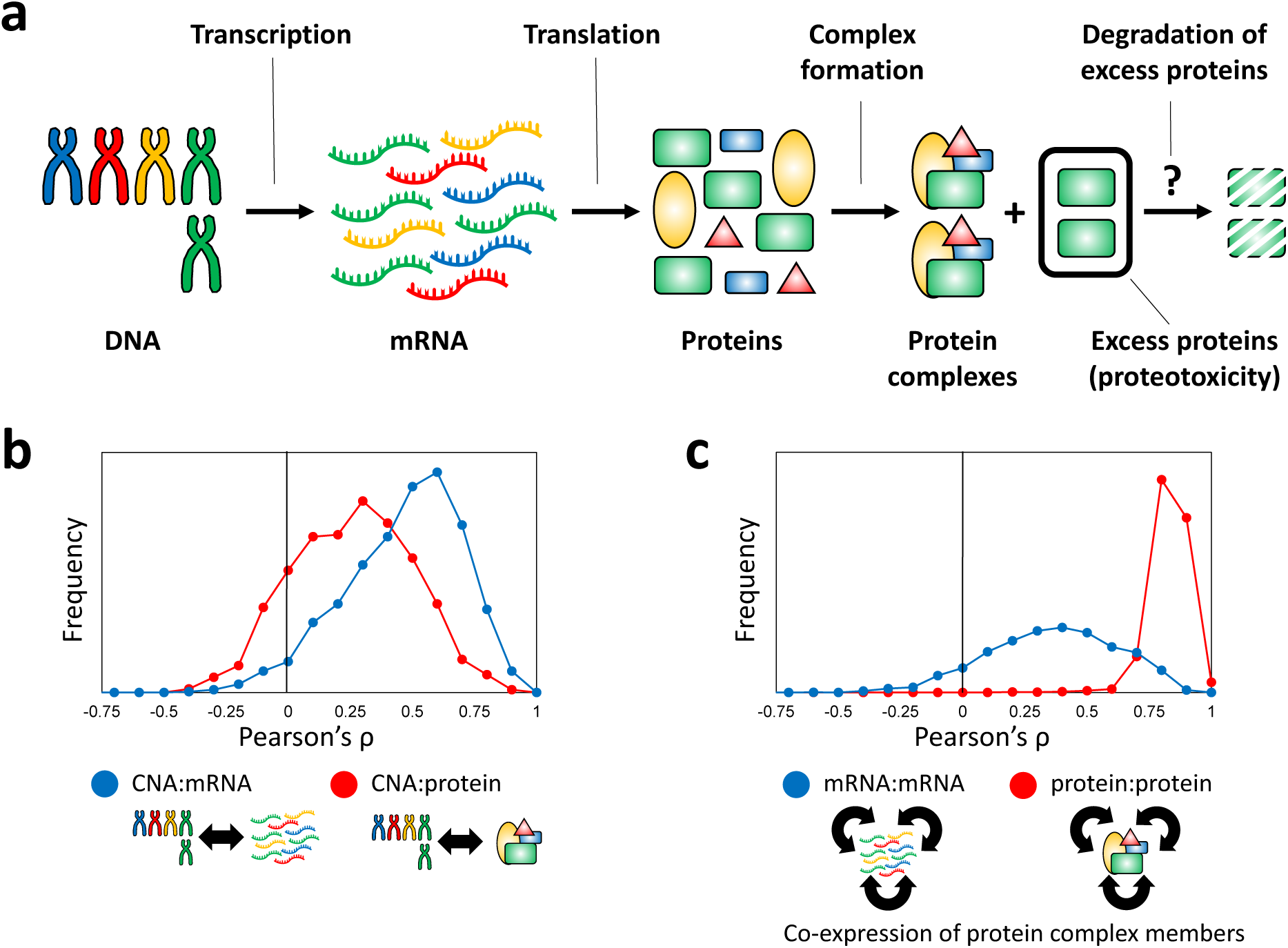
Multi-protein complex protein homeostasis in aneuploid cells. (**a**) Protein homeostasis model for members of multi-protein complexes in aneuploid cells (*30*). In our example, four protein complex members are encoded from different chromosomes, one of which is present with an extra two copies. Gene copy number variation (CNA) levels are maintained through transcription and translation. Targeted degradation of excessive copies of a protein complex member upon complex formation prevents proteotoxicity. This model is supported by a higher correlation of CNA levels with mRNA rather than with protein concentrations (**b**) and by higher co-regulation of protein rather than mRNA concentrations of members of multi-protein complexes (**c**) in human cancer cell lines. Proteomics and RNA sequencing (RNA-seq) data are from a panel of 36 breast cancer cell lines (*11, 13*). Only known members of multi-protein complexes as defined by the CORUM database (*14*) were considered for the data shown in (**b**) (n = 1754, median mRNA:CNV ρ = 0.50, median protein:CNV ρ = 0.27, *P*<10^-15^ (two-sample Kolmogorov-Smirnov test), table S1). For (**c**) associated protein complex members were identified from protein or mRNA co-expression (*11, 13*) (FDR (Benjamini Hochberg) = 5×10^-4^) and filtered for CORUM database confirmed interactions (n (associations) = 2565, median mRNA:mRNA ρ = 0.37, median protein:protein ρ = 0.84, *P*<10^-15^ (two-sample Kolmogorov-Smirnov test), table S1).

We reasoned that the aneuploidy-caused proteotoxicity burden could be defined by the amount of excess protein that must be degraded to level protein concentrations between partners across all protein-protein interactions. To a first approximation, this amount is represented by the absolute CNA difference between two interacting proteins (Fig. 2a), which we have termed copy number imbalance (CNI). To identify the system-wide burden on a cancer cell, we determined a mean CNI value (|CNI|) by averaging CNI values across 13,695 protein-protein interactions identified through protein co-regulation analysis across 41 breast cancer cell lines (Fig. 2a, table S2) (*11*).

**Figure 2.**
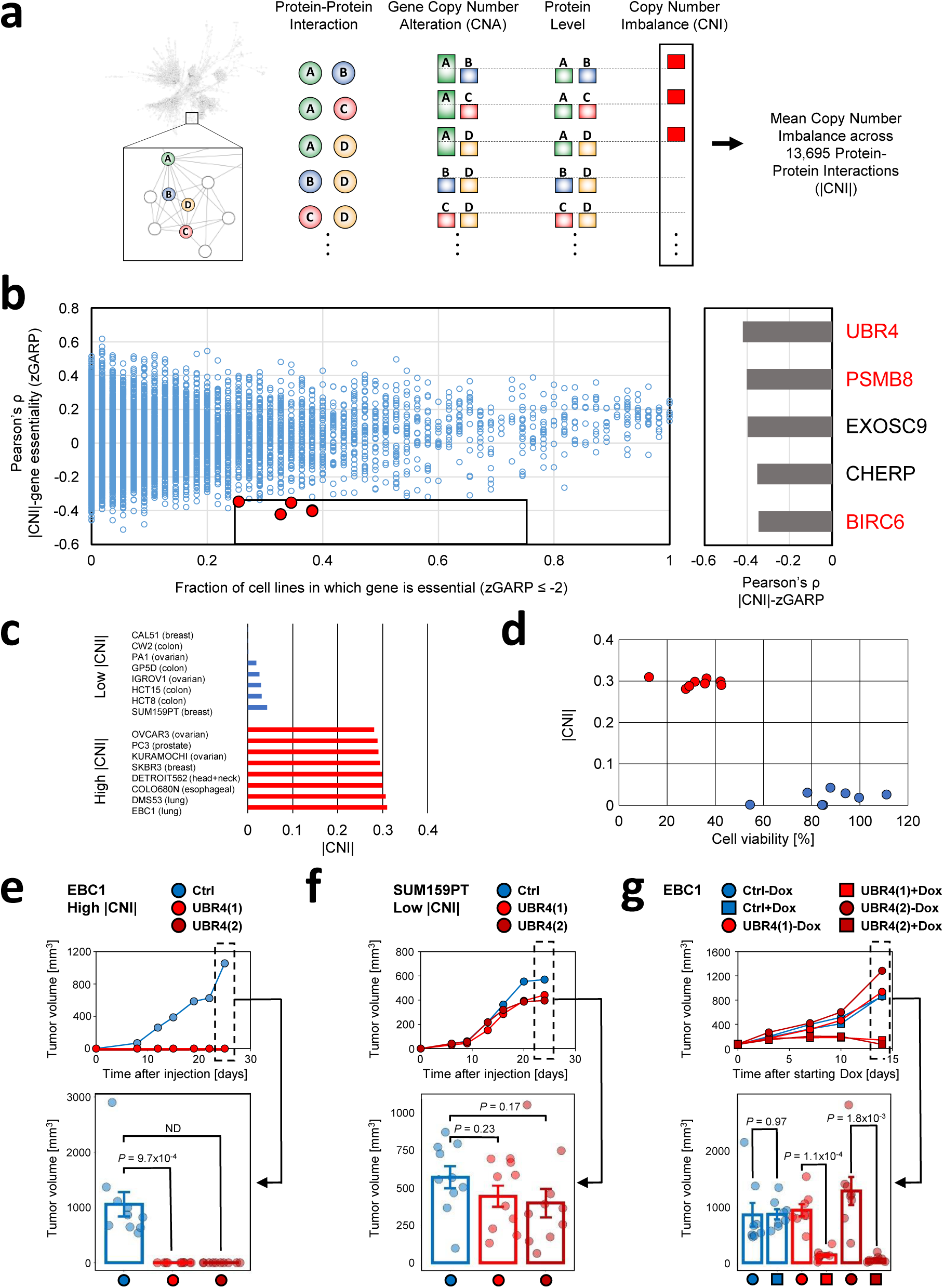
UBR4 is essential for the in vitro and in vivo viability of cancer cell lines with high copy number imbalance of interacting proteins. (**a**) Schematic of the calculation of a cancer’s mean copy number imbalance (|CNI|) defined as the average CNA imbalance for 13,695 protein-protein interactions identified by protein level co-regulation across 41 breast cancer cell lines(*11*). (**b**) For 15697 genes across 55 breast cancer cell lines, the plot shows on the x-axis the fraction of cell lines substantially affected by the depletion of a gene (defined as zGARP ≤ -2, (*16*) versus the correlation of |CNI| and zGARP across all cell lines (y-axis). A group of five genes (red) were found to have a significant negative correlation of |CNI| and zGARP (*P*<0.01, Student’s t-test based on Pearson’s ρ, and zGARP ≤ -2 more than 25 but less than 75 % of the cell lines). Genes with roles in the regulation of protein ubiquitinylation or protein degradation are highlighted in red in the table on the right. PSMB8 and EXOSC9 data points are very similar and almost indistinguishable on the scatter plot (PSMB8, ρ = -0.4, fraction = 0.38; EXOSC9, ρ = -0.39, fraction = 0.38). (**c**) Eight low (blue) and eight high (red) |CNI| cancer cell lines from various cancer types were used for UBR4 depletion experiments. (**d**) The effect of siRNA-enabled depletion of UBR4 on cell viability of low (blue) and high (red) |CNI| cell lines (see fig. S3 and table S4 for depletion efficiency data). Viability is shown relative to control samples. The average viability upon UBR4 depletion was 32 % for high and 87 % for low |CNI| cell lines (*P*=5.6×10^-6^, Student’s t-test). (**e-g**) Effect of UBR4 knockdown on the growth of high and low |CNI| cancer cell lines *in vivo* (xenograft tumors in immunocompromised SCID mice). Subcutaneous tumor growth of a high |CNI| cell line, EBC1(**e,** data for shUBR4 (1) and (2) overlap in the upper graph), and a low |CNI| cell line, SUM159PT (**f**), expressing two independent UBR4 shRNAs(1 and 2) or a scrambled control shRNA(ctrl). The graphs show serial changes in tumor volume (upper), and final tumor volume (lower). (**g**) Growth retardation of already-established EBC1 (high |CNI|) tumors after expression of (Dox)-regulated UBR4 shRNA in vivo. Cell lines harboring Dox-inducible shControl or shUBR4 were injected to generate subcutaneous tumors in SCID animals. Once tumors reached a volume of ∼60-80 mm^3^, Dox was administered in the drinking water. *P* values for f, g, and h, were calculated using the t-test. Depletion efficiency data are presented in fig. S3 and table S4). Data shown in e-g are comprised in table S4.

To identify proteins selectively required for the viability of highly aneuploid cells and potentially involved in adjusting concentrations of interacting proteins with underlying differential CNA levels, we determined the Pearson correlation between |CNI| values and the essentialities of 15,697 genes across 55 breast cancer cell lines (Fig. 2b, table S3; 32 of these cell lines were profiled in our proteomics analysis). The latter were derived from a published pooled genome-wide short hairpin (shRNA) loss-of-function screen, reported in normalized Gene Activity Ranking Profile (zGARP) units, with negative values indicating increased essentiality (only cell lines for which genetic screen and CNA data (*13*) were considered) (*16*). Five genes (UBR4, PSMB8, EXOSC9, CHERP, BIRC6) showed a significant (P<0.01) negative correlation of |CNI| and zGARP while being essential (zGARP ≤ -2) in at least 25 but not more than 75 % of the 55 cell lines, a filter applied to exclude both rare and common essential genes (Fig. 2b and fig. S1). Notably, three of the genes have known roles in the regulation of protein ubiquitinylation or protein degradation (UBR4, PSMB8, BIRC6), and CHERP encodes an endoplasmic reticulum (ER) located protein, consistent with the hypothesis that the unfolded protein response including protein degradation is essential in overcoming aneuploidy-caused proteotoxicity (Fig. 1a).

UBR4 showed the highest correlation between essentiality and |CNI| across the 55 breast cancer cell lines (Fig. 2b). The gene encodes an N-recognin ubiquitin E3 ligase in the N-degron pathway (*17*), a proteolytic system that identifies substrates through protein N-terminal sequences and targets them for degradation via the ubiquitin-proteasome system or autophagy (*18*). UBR4 has also been associated with protein quality control, promoting clearance of misfolded protein aggregates (*19, 20*) and with regulating integrated stress response (*21*). We sought to investigate the essentiality of UBR4 broadly across aneuploid cancers from diverse tissues. To this end, |CNI| values were calculated for 668 cancer cell lines (fig. S2 and table S2) (*13*). We then compared cell viability upon siRNA-induced UBR4 depletion in eight low |CNI| and eight high |CNI| cancer cell lines from either end of the |CNI| distribution. The 16 cell lines models were from breast (n = 3), colon (n = 4), esophageal (n = 1), head and neck (n = 1), lung (n = 2), ovarian (n = 4), and prostate (n = 1) cancers (Fig. 2c). Upon 72 hours of UBR4 depletion, we observed significantly reduced viability in the high |CNI| cell lines (average 32 %) when compared to the low |CNI| cell lines (average 87 %) (*P*=5.6×10^-6^, Student’s t-test) (Fig. 2d, fig. S3 and table S4). Thus, UBR4 is selectively essential for the *in vitro* growth of high |CNI| cell lines derived from a wide range of cancer types. Notably, there was no correlation between relative UBR4 mRNA or protein expression and UBR4 dependence, consistent with a distinct functional role of this protein in the context of high |CNI| (fig. S4).

We next explored the relevance of these findings *in vivo*. To this end, we stably infected the high |CNI| lung cancer cell line EBC1 and the low |CNI| breast cancer cell line SUM159PT with lentiviruses expressing two different shRNAs against UBR4 or with control shRNA. Subcutaneous injection of these cells into immunocompromised SCID mice revealed that EBC1 xenograft tumor growth was completely abrogated upon UBR4 depletion (Fig. 2e, *P*=9.7×10^-4^, Student’s t-test), whereas SUM159PT xenografts were virtually unaffected (Fig. 2f, *P*=0.23) (fig. S3 and table S4). We also assessed the effect of acute depletion of UBR4 in fully formed EBC1 xenograft tumors using a doxycycline-inducible system to express either shRNA targeting UBR4 or control. Doxycycline supplementation was administrated once tumors reached 50 mm^3^. Remarkably, UBR4-depleted tumors showed prominent regression, whereas control-treated tumors continued to grow (Fig. 2g, *P*=1.1×10^-4^, *P*=1.8×10^-3^, two shRNA probes were used, fig. S3 and table S4). Thus, UBR4 is required for both xenograft tumor formation and maintenance in the context high of high |CNI| cancer.

We sought to gain insight into the biological role of UBR4 in aneuploid cancer by comparison of the differential protein expression changes caused by UBR4 inactivation in high versus low|CNI| cell lines. In particular, we predicted that this analysis would highlight a distinct cellular response program resulting from UBR4 depletion and also reveal a convergence of CNA and protein concentrations as an underlying mechanism. Accordingly, we quantitatively mapped proteome changes in the 16 low and high |CNI| cell lines upon shRNA-induced UBR4 depletion using two different shRNA probes versus control shRNA. We applied multiplexed quantitative proteomics using eleven-plex tandem mass tag (TMT) reagents and the SPS-MS3 method on an Orbitrap Lumos mass spectrometer to achieve high quantitative accuracy (*22–25*). Samples were analyzed in duplicates to generate 96 proteome maps (Fig. 3a). A total of 10,796 proteins were quantified across all samples, and 7,187 proteins were quantified in all cell lines and conditions (upon combining data from duplicate measurements, fig. S5 and table S5). We characterized the proteins differentially upregulated upon UBR4 depletion in high versus low |CNI| cells by first calculating protein ratios for shUBR4 over shControl samples for each cell line, and then by averaging the ratio of these ratios for high over low |CNI| cell lines (HCT15 was excluded from the analysis based on an insufficient knockdown of UBR4, fig. S5). To gain a global view of the pathway alterations caused by UBR4 depletion, we ranked the proteins according to their t-statistics and analyzed them using Gene Set Enrichment Analysis (GSEA) (*26*), first considering only cell compartment categories. Leading-edge genes from categories found to be significantly (FDR < 15 %, Kolmogorov-Smirnov test) upregulated in UBR4-depleted high |CNI| cells were clustered into broader categories using the DAVID bioinformatics platform (*27*). Significantly upregulated clusters (EASE score > 10, Benjamini Hochberg FDR≤0.25, Fisher’s Exact test) are shown in Fig. 3b (table S6). The highest scoring cluster consisted of ER resident proteins, which—together with enrichment of other clusters of proteins assigned to the lysosome, vesicles, and vacuoles –was in keeping with our hypothesis that UBR4 is essential to the unfolded protein response in high |CNI| cells that have elevated concentrations of misfolded proteins. This was further supported by analyzing the ER-located proteins for enrichment of functional categories of biological processes, which revealed response to ER stress as the most significantly upregulated pathway and demonstrated prominent increases in response to topologically incorrect and unfolded proteins (Fig. 3c, FDR≥1×10^-6^, table S6). The relative expression changes of proteins in networks that correspond to these latter two clusters are shown in Fig. 3d. No categories were significantly upregulated in UBR4-depleted low |CNI| cell lines using the same analytical strategy (table S6). Therefore, UBR4 inactivation causes evidence ER stress and unfolded protein response, specifically in high |CNI| cancer cell lines.

**Figure 3.**
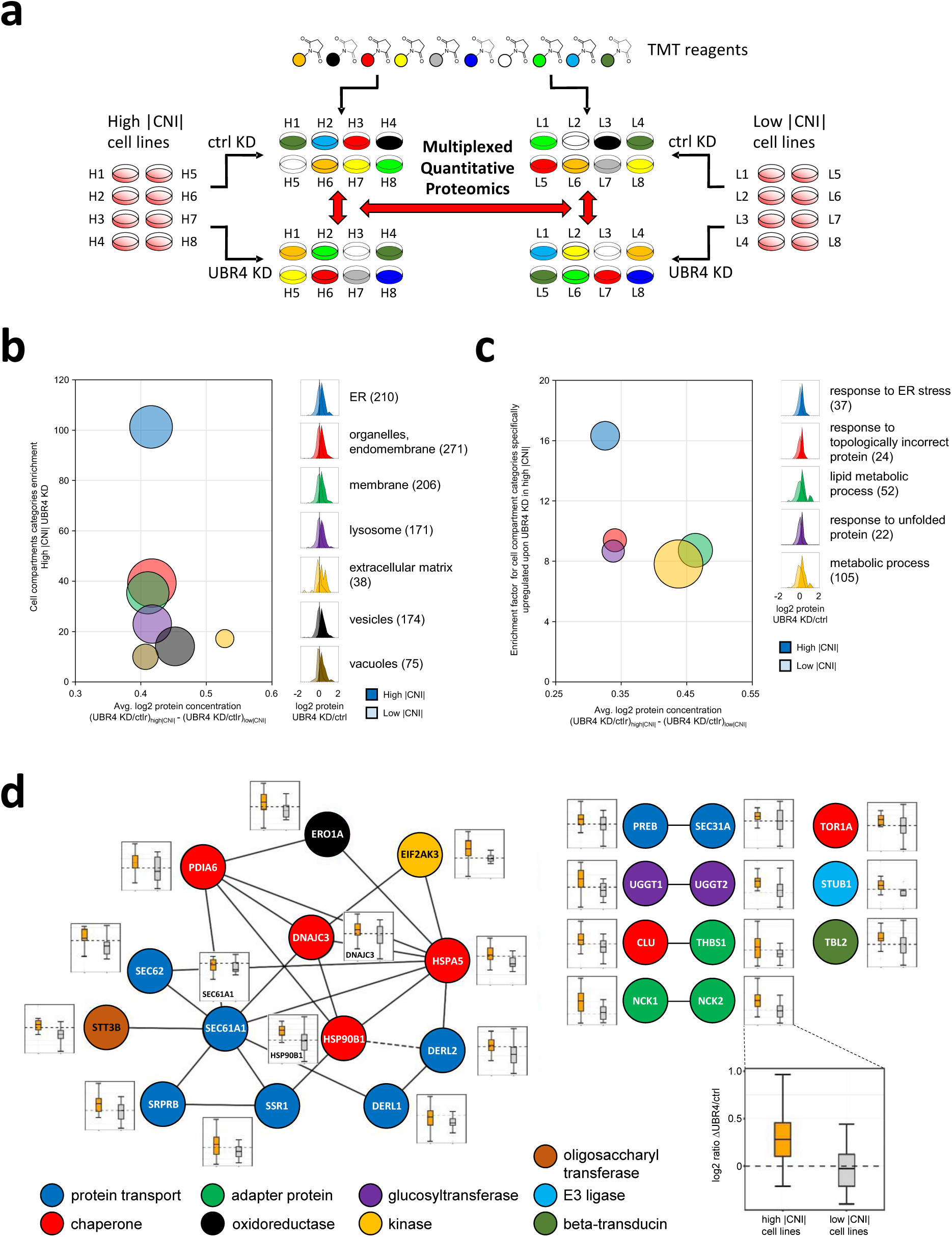
UBR4 depletion in high |CNI| cancer cells increases concentration of proteins regulating protein quality control. (**a**) Multiplexed quantitative mass spectrometry-based proteomics workflow using tandem mass tag (TMT11) technology. Proteomes were mapped in duplicate for eight high |CNI and eight low |CNI| cell lines (see Fig 2d) upon shRNA-based depletion of UBR4 using two UBR4 targeting shRNA probes and one control shRNA (see Fig 2f and 2g). A total of 96 samples were analyzed, 10796 proteins were quantified across all samples, and 7187 in all conditions. Data from the cell line HCT15 were not considered for further analysis due to an insufficient UBR4 depletion (table S5 and fig. S5). (**b**) ‘Cellular component’ (CC) gene ontology categories (*27*) enriched in proteins significantly upregulated in high |CNI| but not in low |CNI| cell lines upon UBR4 depletion compared with ctrl depletions [table S6; circle size represents the number of genes in each category clusters (numbers are also listed in parentheses on the right); histograms on the right show the differential regulation of the protein categories in high and low |CNI| cell lines; see Methods for data analysis details]. (**c**) ‘Biological process’ (BP) gene ontology categories enriched in proteins contained in the CC component ‘Endoplastmic Reticulum’ (ER) shown in (b) (table S6). (**d**) Network of proteins included in the two components ‘response to topologically incorrect protein’ and ‘response to unfolded protein’ categories shown in (c). Protein interactions were extracted from the STRING database (*31*) (high evidence, solid line) or determined through protein co-regulation analysis (*11*) (dotted line). Inserted plots show the log2 protein ratio in UBR4 depleted samples (averaged data from two shRNA probes) over ctrl samples in high (orange) and low (grey) |CNI| cell lines.

Upon showing the essentiality of UBR4 for the viability of cancer cells with a large CNA burden, and its association with unfolded protein response, we examined our prediction that UBR4 levels the concentrations of proteins in multi-protein complexes to suppress proteotoxicity. In accord with such a role, we expected that UBR4 depletion would reduce co-regulation of the concentration of complex members in high |CNI| cells leading to convergence of CNA and protein levels. Previously, we described measuring deviations in the co-regulation of protein pairs in canonical complexes as a means to identify cell line-specific dysregulations of protein-protein interactions (*11*). Such deviations – determined by measuring Mahalanobis distances (MD) for each cell line in pairwise protein concentration comparisons – indicate a perturbation in the adjustment of protein levels of the affected protein-protein interactions. We reasoned that UBR4 depletion in high |CNI| cell lines should cause a global increase in these deviations. Our analytical workflow to test this hypothesis is shown in Fig. 4a: proteome data from 41 breast cancer cell lines was used to generate an interactome dataset of 16747 protein-protein interactions, which subsequently served as a background-network to identify UBR4 depletion-induced deviations of protein concentration co-regulation in low and high |CNI| cell lines (see Methods, HCT15 was excluded based on insufficient UBR4, knockdown fig. S5). An example of a deviated co-regulation between HEATR1 and RSL1D1 upon UBR4 depletion in the ovarian cancer cell line OVCAR3 is shown in Fig. 4b (more examples – one for each high |CNI| cell line – are shown in fig. S6). Defining a deviation in protein co-regulation induced by UBR4 depletion as a two-fold increased MD over the shControl sample (average of two independent shRNAs against UBR4, coefficient of variation (CV) ≤ 0.25, Methods), we determined the number of deviations for the 15 cancer cell lines (Fig. 4c). Strikingly, high |CNI| cell lines dominated the list of cancer cell lines with the greatest number of induced deviations, except for HCT8 and SUM159PT, which were categorized as low |CNI| cell lines but had highest |CNI| values within this group (Fig. 2c) (table S7). The Spearman correlation coefficient between the number of induced co-regulation deviations and |CNI| values was 0.46 (*P*=0.088) indicating a strong trend of a higher number of deviations in high |CNI| cell lines and confirming our hypothesis on the role of UBR4 in adjusting protein levels.

**Figure 4.**
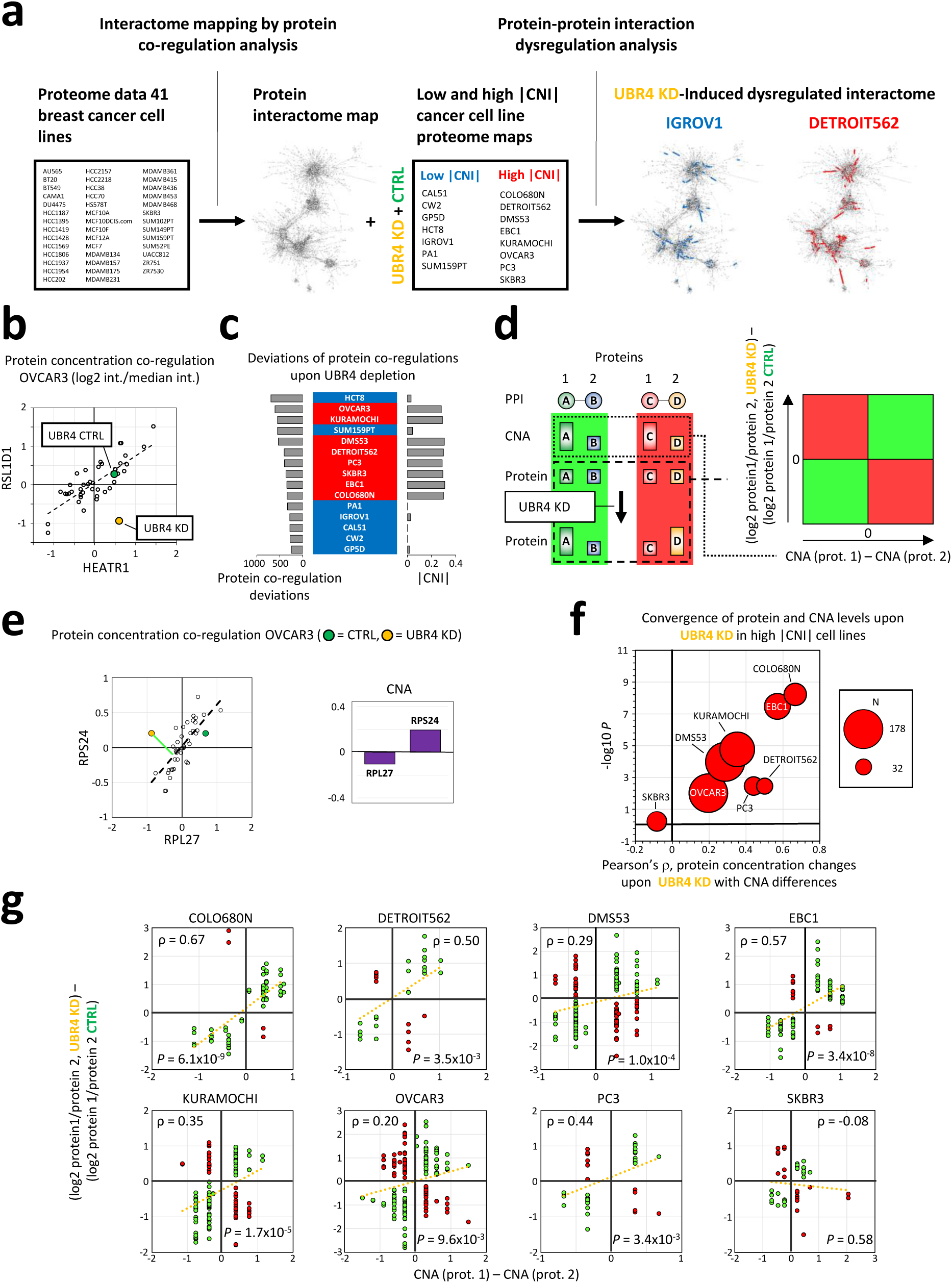
UBR4 depletion disrupts protein concentration co-regulation of interacting proteins and aligns protein with CNA levels. (**a**) Analytical workflow to determine the role of UBR4 in the co-regulation of the concentration of interacting proteins. A protein co-regulation map (16747 interactions, table S7) was generated using proteome data from 41 breast cancer cell lines only considering proteins quantified in all cell lines (breast cancer and the current study, see Methods) (*11*). UBR4 depletion-induced deviations from the identified co-regulations were determined based on the Mahalanobis distance (MD) for seven low |CNI| and eight high |CNI| cell lines upon UBR4 depletion (average MD in two UBR4 KD samples ≥ 2-fold over ctrl sample, MD coefficient of variance (CV) in the two KD ≤ 0.25, table S7). (**b**) An example of a UBR4 KD-induced co-regulation deviation for the protein pair HEATR1 and RSL1D1 in the OVCAR3 cell line. The other points plotted represent levels of these proteins at baseline across the collection of 41 breast cancer cell lines. (**c**) Protein coregulations upon UBR4 depletion in the studied 15 cancer cell lines. High |CNI| cell lines are shown in red, low |CNI| cell lines in blue. The number of induced deviations is shown in the bar plot on the right, and the |CNI| values of the cell lines on the left. The Spearman correlation factor between induced deviations and |CNI| is 0.46 (*P* = 0.088) (**d**) Analytical workflow to determine correlations between protein concentration changes underlying deviations of protein co-regulations upon UBR4 depletion and CNA levels of encoding genes. Green indicates that the protein and CNA levels are converging, red indicates an increasing difference between the levels. When measurements for all interaction deviations for a cell line are shown in scatter plot (right), a positive significant Pearson correlation indicates convergence across all data points. (**e**) The protein concentrations of RPL27 and RPS24 converge with CNA levels upon UBR4 depletion in OVCAR3 (protein and CNA levels are log2 ratios over the median signal of all protein and CNA levels, respectively). (**f**) Monitoring convergence of protein and CNA levels in protein-protein interactions upon UBR4 depletion in eight high |CNI| cell lines. The analysis was done as described in (d). Circle size reflects the number of protein co-regulation deviations analyzed for each cell line. In addition to a 2-fold change of the MD upon UBR4 depletion (a), we required deviations to be significant (MD-based bivariate outlier testing (Grubb’s test, FDR=25 %, table S8, Methods). (**g**) Monitoring convergence of protein and CNA levels in protein-protein interactions upon UBR4 depletion in each individual high |CNI| cell line. All cell lines but SKBR3 showed a significant convergence.

Next, we tested whether the deviations from protein co-regulation caused by UBR4 depletion in high |CNI| cell lines involved a convergence of protein and CNA levels. The analytical concept of this analysis is shown in Fig. 4d. Concentrations of co-regulated protein pairs within a canonical complex may undergo a convergence with CNA levels upon UBR4 depletion (Fig. 4d, green areas) or change in a manner to increase the difference between protein and CNA levels (red areas). For multiple deviations in co-regulation, these relationships can be shown by plotting changes in the ratio of the concentrations of each protein pair versus the underlying CNA ratios (Fig. 4d, right; green quadrants show convergence, red quadrants show an increased gap between protein and CNA levels; a positive correlation across all data levels upon UBR4 depletion induced co-regulation deviation indicates an overall convergence). Fig. 4e shows an example of the convergence of protein and CNA levels for the protein pair RPL27 and RPS24 in the OVCAR3 cell line (more examples – one for each high |CNI| cell line – are shown in fig. S6). Across all significant co-regulation deviations [bivariate MD-based Grubb’s outlier test (FDR=25 %)] shown in Fig. 4c and table S8 (Methods), we found a significant convergence of protein level changes upon UBR4 depletion and CNA levels across all high |CNI| cell lines except SKBR3 (Fig. 4f, overview plot; Fig. 4g, individual cell lines). Taken together, these results strongly suggest that UBR4 is a key regulator of global protein homeostasis, with specific roles in adjusting concentrations of canonically interacting proteins.

Here we used integrative analysis of proteomics and genome-wide loss-of-function screening data to identify the E3 ligase UBR4 as a key component in the accurate adjustment of protein concentrations within multi-protein complexes. By preventing the proteotoxic accumulation of excess proteins that exceed the stoichiometric needs of their binding partners, our data suggest that UBR4 – and by extension, the N-degron pathway – plays a critical role in shaping the cellular interactome. Conceptually, these findings imply that protein degradation by the N-degron pathway may provide a critical tension between protein folding and the formation of quaternary structures, where N-terminal degrons may be buried within assembled complexes, while those subunits that cannot find their binding partners are being degraded before they can cause toxicity.

Our data show that UBR4 plays an instrumental role in cancer by overcoming the CNA-derived proteotoxicity that constitutes the collateral damage of aneuploidy. Since aneuploidy is extremely widespread in cancer, UBR4 may present a target for a broadly applicable cancer treatment strategy. Mouse knockout studies show that UBR4 is required for embryogenesis (*17*), and mutations may play a role in degenerative neurological conditions (*28*). However, we think that its lack of essentiality for non-aneuploid cancer cells – as shown in this study – may open a therapeutic window for using UBR4 inhibitors in disease treatments. Importantly, aneuploidy is not only observed in cancer but may also play a role in aging (*29*). UBR4 inhibition may, therefore, present a therapeutic strategy extending to other aging-associated diseases.

## Acknowledgments

We thank Steven Gygi, Harvard Medical School, for access to computational software and facilities to process the proteomics data. We acknowledge all members of the Haas and Bardeesy laboratories for valuable discussions.

## Data and materials availability

All data are available in the main text or the supplementary materials. All mass spectrometry RAW data will be made accessible through the MassIVE data repository (massive.ucsd.edu). Knockdown cell lines will be provided upon request.

## Supplementary Materials

### Materials and Methods

#### Tissue culture

Cancer cell lines were grown under the indicated conditions (table S4). Adherent cells were rinsed with PBS before being trypsinized and were counted and seeded at appropriate concentrations for further experiments. For western and MS analysis, cells were pelleted at 1000 rpm for 5 min at 4 °C. Pellets were washed 2X in PBS before being fast-frozen on dry ice then stored at -80 °C until lysis.

#### *In vitro* cell viability assay

Transient knockdown of UBR4 was accomplished by reverse transfection using SMARTpool siRNAs (Dharmacon GE, Pittsburgh, PA, USA). A final siRNA concentration of 10 nM and after 72 h a second forward transfection with 20 nM siRNA was delivered to the cells with *Lipofectamine 3000* Transfection Reagent according to the manufacturer’s protocol. The transfection was performed in clear bottom 96-well plates from Corning. Sixteen cancer cell lines were treated with a SMARTpool comprised of 4 different siRNAs targeting UBR4, an ON-TARGET siRNA targeting GFP, 4 ON-TARGET non targeting siRNAs, and a SMARTpool comprised of 4 different non-targeting siRNAs. Cells were incubated with siRNA for 144 h and cell viability was evaluated by measuring intracellular ATP concentrations using the *CellTiter*-*Glo* luminescent cell viability assay (Promega Corp., Madison, WI, USA). Chemical luminescence was measured using a microplate reader and knockdown efficiency was verified by Western blotting.

#### Preparation and transduction of lentiviral-delivered short-hairpin RNA (shRNA)

For stable knockdown of UBR4 expression, pLKO.1-Puro lentiviral constructs expressing shRNA against human UBR4 were obtained from the MGH Molecular profiling laboratory (MPL) at the MGH Cancer Center. The UBR4 mRNA sequences targeted with lentiviral shRNA clones are, [5’- CGAGCCATTCTACTCATCTTT-3’, TRCN0000155927, labeled as A5 (probe 1)] and [5’- GCCGACTAGATAGAACTGAAA-3’, TRCN0000155202 labeled as C1 (probe 2)]. pLKO.1- scrambled shRNA (Addgene Plasmid repository ID #1864, originally created by David M. Sabatini, MIT) was used as a control. For doxycycline-inducible knockdown of UBR4, A5 (shUBR4 1) and C1 (shUBR4 2) shRNA sequences were cloned in pLKO-Tet-On-shRNA lentiviral vectors (Addgene Plasmid repository ID # #21915, originally created by Dmitri Wiederschain, Novartis). A Scrambled shRNA(5’- CCTAAGGTTAAGTCGCCCTCG-3’) cloned in pLKO-Tet-On was used as control. Lentiviral particles were generated in 293T cells with pCMV-dR8.2 dvpr and pCMV-VSV-G lentiviral packaging plasmids with XtremeGene 9 Transfection Reagent (Sigma). The spent medium containing lentiviral particles was collected, filtered through a 0.45 μm syringe filter (Fisher Scientific), and stored at -80°C until further use. Tumor cells were infected with viral supernatant and polybrene (8 μg/ml) for 48 hrs. Then cells were selected for 72 hrs in the presence of puromycin (2 μg/ml) to establish lines expressing shRNA constructs. For the dox-inducible system, 100-1000ng/ml doxycycline was used to verify the efficiency of knockdown of UBR4.

#### Animal studies

Mice were housed in pathogen-free animal facilities. All experiments were conducted under protocol 2005N000148 approved by the Subcommittee on Research Animal Care at Massachusetts General Hospital. Data presented include both male and female mice. All mice included in the survival analysis were euthanized when criteria for disease burden were reached (maximum approved tumor size, loss of >15% of body weight, compromised ambulatory function, labored breathing, and/or abnormal posture). For subcutaneous tumor studies, 1.0-1.5×10^6^ shRNA modified cells were subcutaneously injected into the lower flanks of 6-10 weeks old NOD.CB17-Prkdcscid/J mice (Jackson Laboratories, strain #001303). Tumor size was assessed at indicated time points by caliper measurements of length and width and the volume was calculated according to the formula ([length × width2]/2). For dox inducible shRNA experiments tumor cells expressing pLKO-Tet-On scrambled, C1 and A5 shRNAs were injected subcutaneously and allowed to form tumors. Once the tumor attained an average size of ∼60-80 mm^3^ diameter, the animals from each shRNA group were randomly divided into two cohorts. One cohort received dox (200 μg/ ml) in drinking water to express shRNAs, and the other received normal water serving as the control cohort. Tumor size was measured as described above.

#### Proteomics

##### Cell lysis, protein digestion, and TMT labeling

Cell pellets were lysed with 600 µL of lysis buffer by passing through a 21-gauge needle 20 times. Lysis buffer was composed of 75 mM NaCl, 3 % SDS, 1 mM NaF, 1 mM beta-glycerophosphate, 1 mM sodium orthovanadate, 10 mM sodium pyrophosphate, 1 mM PMSF and 1X Roche Complete Mini EDTA free protease inhibitors in 50 mM HEPES, pH 8.5. Proteins were then reduced with DTT at 56 ° C for 30 min and alkylated with iodoacetamide as previously described (*25*). Tricholoracetic acid (TCA) was used to precipitate the reduced and alkylated proteins and dried pellets were reconstituted in 300 μL of 1 M urea in 50 mM HEPES, pH 8.5. Proteins were solubilized using a combination of vortexing, sonication, and manual grinding and then digested in a two-step process starting with overnight digestion at room temperature with 3 μg of Lys-C (Wako) followed by six hours of digestion with 3 μg of trypsin (sequencing grade, Promega) at 37 °C. The digest was acidified with 10 % TFA and peptides were desalted on C_18_ solid-phase extraction (SPE) (Sep-Pak, Waters) columns as previously described (*25*). The concentration of the desalted peptide solutions was measured with a BCA assay, and peptides were aliquoted into 50 μg portions, dried under vacuum, and stored at -80 °C until they were labeled with TMT reagents (*25, 32*).

TMT labeling was performed with 11-plex tandem mass tag (TMT) reagents (Thermo Scientific) as previously described (*25*). TMT reagents were dissolved in dry acetonitrile (ACN) at a concentration of 20 μg/μL. Dried peptides (50 μg) were re-suspended in 30 % dry ACN in 200 mM HEPES, pH 8.5 and 4.5 μL of the appropriate TMT reagent was added to the sample. TMT reagent 126 was reserved for “bridge” samples which is a pooled sample composed of all samples including replicates (*11*). The remaining TMT reagents (127N, 127C, 128N, 128C, 129N, 129C, 130N, 130C, 131N, 131C) were used to label the digests from the individual cell lines in a random order (see table S5 for the TMT labeling scheme). Peptides were incubated with the reagents for 1 h at room temperature. The labeling reaction was quenched by adding 5 μL of 5 % hydroxylamine. Labeled samples were then acidified by adding 30 μL of 1 % TFA and the peptide mixtures were pooled into 10 11-plex TMT samples, with the bridge sample carrying the 126 label. The pooled samples were desalted via C_18_ SPE on Sep-Pak cartridges as described above.

##### Basic pH reversed-phase liquid chromatography (bRPLC) sample fractionation

The 11-plex TMT-labeled samples were fractionated by basic pH reversed-phase liquid chromatography (bRPLC) with concatenated fraction combining as previously described (*25*). Briefly, samples were re-suspended in 5 % formic acid/5 % ACN and separated over a 4.6 mm x 250 mm ZORBAX Extend C_18_ column (5 μm, 80 Å, Agilent Technologies) on an Agilent 1260 HPLC system outfitted with a fraction collector. The separation was performed by applying a gradient build from 22 to 35 % ACN in 10 mM ammonium bicarbonate in 60 minutes at a flowrate of 0.5 mL/minute. A total of 96 fractions were combined as previously described. The combined fractions were dried under vacuum, re-constituted with 5 % formic acid/5 % ACN, and analyzed by LC-MS2/MS3 for identification and quantification.

##### Liquid chromatography mass spectrometry

All LC-MS2/MS3 experiments were conducted on an Orbitrap Fusion Lumos mass spectrometer (Thermo Fisher Scientific) coupled to an Easy-nLC 1200 (Thermo Fisher Scientific) with a chilled autosampler. Peptides were separated on an in-house pulled, in-house packed microcapillary column (inner diameter, 100 μm; outer diameter, 360 μm). Columns were packed first with approximately 0.5 cm of Magic C_4_ resin (5 μm, 100 Å, Michrom Bioresources) followed by approximately 0.5 cm of Maccel C_18_ AQ resin (3 μm, 200 Å, Nest Group), and then to a final length of 30 cm with GP-C_18_ (1.8 μm, 120 Å, Sepax Technologies). Peptides were eluted with a linear gradient from 11 to 30 % ACN in 0.125 % formic acid over 165 minutes at a flow rate of 300 nL/minute while the column was heated to 60 °C. Electrospray ionization was achieved by applying 1800 V through a stainless-steel T-junction at the inlet of the microcapillary column.

The Orbitrap Fusion Lumos was operated in data-dependent mode, with a survey scan performed over an m/z range of 500-1,200 at a resolution of 6×10^4^ in the Orbitrap. For the MS1 survey scan, automatic gain control (AGC) was set to 5 x 10^5^ and a maximum injection time of 100 ms. The S-lens was set to an RF of 30 and data was centroided. The most abundant ions detected in the survey scan were subjected to MS2 and MS3 experiments using the Top Speed setting that enables a maximum number of spectra to be acquired in a 5 seconds experimental cycle before the next cycle is initiated with another survey full-MS scan.

For MS2 analysis, the decision tree option was enabled, whereby precursors were selected based on charge state and m/z range. Doubly charged ions were selected from an m/z range of 600-1200, for triply and quadruply charged ions had to be detected in an m/z range of 500-1200. The ion intensity threshold was set to 5×10^4^. When acquiring MS2 spectra ions were isolated by applying a 0.5 m/z window using the quadrupole and fragmented using CID at a normalized collision energy of 30 %. Fragment ions were detected in the ion trap at a rapid scan rate. The AGC target was set to 2 x 10^4^ and the maximum ion injection time was 35 ms. Centroided data was collected. MS3 analysis was performed using synchronous precursor selection (MultiNotch MS3) enabled to maximize sensitivity for quantification of TMT reporter ions (*22, 24*). Up to five MS2 precursors were simultaneously isolated and fragmented for MS3 analysis. The isolation window was set to 2 m/z and fragmentation was carried out by HCD at a normalized collision energy of 55 %. Fragment ions in the MS3 spectra were detected in the Orbitrap at a resolution of 5 x 10^4^ at an m/z of ≥ 110. The AGC target was set to 4.5 x 10^5^ ions and the maximum ion injection time to 150 ms. Centroided data were collected. Fragment ions in the MS2 spectra with an m/z of 40 m/z below and 15 m/z above the precursor m/z were excluded from being selected for MS3 analysis.

##### Data processing and analysis

Data were processed using an in-house developed software suite^9^. RAW files were converted into the mzXML format using a modified version of ReAdW.exe (http://www.ionsource.com/functional_reviews/readw/t2x_update_readw.htm). Spectral assignments of MS2 data were made using the Sequest algorithm (*33*) to search the Uniprot database of human protein sequences including known contaminants such as trypsin. The database was appended to include a decoy database consisting of all protein sequences in reverse order (*34, 35*). Searches were performed with a 50 ppm precursor mass tolerance. Static modifications included ten-plex TMT tags on lysine residues and peptide n-termini (+229.162932 Da) and carbamidomethylation of cysteines (+57.02146 Da). Oxidation of methionine (+15.99492 Da) was included as a variable modification. Data were filtered to a peptide and protein false discovery rate of less than 1 % using the target-decoy search strategy (*35*). This was achieved by first applying a linear discriminator analysis to filter peptide annotations (peptide-spectral matches) using a combined score from the following peptide and spectral properties: XCorr, ΔCn, missed tryptic cleavages, peptide mass accuracy, and peptide length (*36*). The probability of a peptide-spectral match to be correct was calculated using a posterior error histogram and the probabilities of all peptides assigned to one specific protein were combined through multiplication and the dataset was re-filtered to a protein assignment FDR of less than 1 % (*36*) for the entire dataset of all proteins identified across all analyzed samples. Peptides that matched to more than one protein were assigned to that protein containing the largest number of matched redundant peptide sequences following the law of parsimony (*36*).

For quantitative analysis TMT reporter ion intensities were extracted from the MS3 spectra selecting the most intense ion within a 0.003 m/z window centered at the predicted m/z value for each reporter ion and a signal-to-noise (S/N) values were extracted from the RAW files. Spectra were used for quantification if the sum of the S/N values of all reporter ions was ≥ 40 times the number of TMT channels, and the isolation specificity for the precursor ion was ≥ 0.75 (*22*). Protein intensities were calculated by summing the TMT reporter ions for all peptides assigned to a protein. Normalization of the quantitative data followed a multi-step process. Intensities were first normalized using the intensity measured for the bridge sample. The average median bridge channel intensity measured across all 11 TMT experiments was used for the normalization. Taking account of slightly different protein amounts analyzed in each TMT channels we then added an additional normalization step by normalizing the protein intensities measured for each sample by the median of the median protein intensities measured in these samples. The proteome profiles from the analyses of two biological replicates were combined by calculating the average intensity if the protein was quantified in both replicates but also including intensities of proteins that were only quantified for one replicate. If the Pearson correlation coefficient between two replicate data sets from the same cell line and condition was <0.5, the dataset with the least average correlation coefficient with the other datasets (all conditions and replicates) of the same cell line was removed from the dataset (only proteins quantified in all replicates were considered for this analysis, normalized intensities were converted into log2 ratios of the intensities over the median intensity measured for each protein across all cell lines). Based on this quality control analysis, replicate 2 of the CAL51 control sample, replicate 2 of the CW2 control sample, and replicate 1 of the OVCAR3 UBR4 KD with shRNA probe 1 were not considered when averaging replicate datasets.

All mass spectrometry RAW data will be accessible through the MassIVE data repository (massive.ucsd.edu).

#### Bioinformatics

##### Relationship between protein, mRNA, and copy number alterations (CNA)

Full proteome profile data for 41 breast cell lines were generated in-house (*11*). Data for mRNA levels and copy number alterations (CNA) were taken from Kljin et al. (*13*). The proteomics profiles were converted into a relative scale by dividing each individual protein level by the median of the protein level across all samples, followed by the log2 transformation of the ratios. The same log2-over-median-intensity transformation was applied to the mRNA profiles. The CNA data was provided as a log2 ratio relative to a diploid state (negative = loss, positive = gain). Proteome, mRNA, and CNA data were available for 36 breast cell lines and enabled generating a combined dataset with no missing values across all three sub-datasets in all cell lines for 5625 genes and gene products. Information on known human protein complexes was extracted from the CORUM database (v10.5) (*14*) and the dataset was further reduced by only considering known as members of human protein complexes (n = 1754; 2565 interactions). Across all cell lines, Pearson correlation coefficients were calculated between protein, mRNA, and CNA levels. In addition, Pearson correlations between protein levels as well as mRNA levels of proteins associated with CORUM-annotated interactions were calculated. Statistical comparisons of correlation coefficients were performed using the Kolomogorov-Smirnov test. Data are listed in table S1.

##### Copy number imbalance (CNI) and gene essentialities

Copy number imbalance (CNI) is a metric representing the inherent effort a cell needs to apply to maintain homeostatic protein levels in protein-protein interactions. The CNI value of an interaction was calculated by taking the difference between the CNA levels of the interacting protein partners (protein_1 CNA - protein_2 CNA). Using CNA data from Kljin et al. (*13*), CNI values were determined for 668 cell lines 13695 protein associations previously identified by protein co-regulation analysis (associations were considered only if CNA data were available) (*11*). Differences between CNA values were averaged across all interactions to calculate the mean CNI (|CNI|) for each cell line (table S2). Utilizing previously published shRNA technology-based genome-wide drop-out screen data (*16*) a set of 55 breast cell lines was identified, for which both, |CNI| values and essentiality data for 15697 genes, were available. Gene essentialities were reported in normalized Gene Activity Ranking Profile (zGARP) units with negative values indicating a reduction of cell fitness. Genes with increased essentiality in high |CNI| cell lines were identified by calculating Pearson correlations between |CNI| and gene essentiality for each gene across the 55 cell lines. Candidate genes were essential (zGARP ≤ -2) in more than 25 % and less than 75 % of the 55 cell lines and showed a significant (P<0.01) negative correlation between zGARP and |CNI|.

##### Differential expression of proteins upon UBR4 depletion

The UBR4 KD induced protein changes were compared between the high and low |CNI| cell lines. The duplicate proteomic profiles for the two knockdown and control samples were averaged to collapse across replicates. Log2 fold changes (shRNA / control) were generated for each of the shRNAs probes (1 and 2). These shRNA log2 fold changes were compared between the eight high |CNI| and seven low |CNI| cell lines for each of the 7187 proteins quantified across all conditions using a Student’s T-test. P-values were adjusted using the Benjamini-Hochberg multiple hypothesis correction.

Functional enrichment analysis: The functional enrichment analysis of the differentially expressed proteins induced by the UBR4 KD across eight high and seven low |CNI| cell lines were performed using the BROAD GSEA tool (*26*). The input for the GSEA tool was an RNK formatted table of protein names converted to unique gene symbols, pre-ranked in descending order based on the T-test statistic. GSEA searched for the enrichment of MSigDB v6.0 (*37*) cellular component GO term genesets using the following command line parameters (-nperm 10,000, -scoring_scheme weighted, -mode Max_probe, -norm meandiv, -setLiberzon et al. 2011_min 15, -set_max 2000). A total of 11 significant cellular component genesets (FDR<0.15) were identified as up-regulated in the KD of the high |CNI| cell line context. A total of 307 genes were identified as leading edge or core enriched genes across the 11 enriched terms. After filtering down to the 307 genes the functional enrichment web tool, DAVID (*38*) was used to look for cellular component enrichment clusters within the set of leading edge genes. The input for the tool was the 307 gene symbols and the background for the analysis was the entrez gene identifiers for the 7187 proteins quantified in the knock-down experiment. The gene ontology GO_TERM_CC_ALL was the only gene set collection selected for the enrichment and we used the medium (default) classification stringency parameters. The DAVID cluster tool sorted the groups of enriched annotation terms based on their cluster enrichment score (EASE), a modified Fisher’s exact p-value. This analysis yielded seven clusters of functional terms with an enrichment score greater than 10. The top cluster was associated with the endoplasmic reticulum. A functional enrichment analysis of the 210 genes associated with the terms in the ER cluster was performed with DAVID using the medium classification stringency parameters and the GO biological process (GO_TERM_BP_ALL) gene set collection. A total of nine GO biological process terms were found to be significant, with an EASE score greater than 6. Four of the nine terms were found to be redundant and removed from the downstream analysis. A redundant term had all member genes encapsulated within another related significant term.

##### Mapping dysregulation protein-protein interactions and convergence or divergence of protein and CNA levels upon UBR4 depletion

Proteome data acquired for the 16 cancer cell lines were combined with the 41 breast cancer cell line protein data using an average normalization methodology. First, the UBR4 knockdown and breast cancer cell line protein profiles were reduced the common set of 5825 proteins. The average normalization was then performed in two steps: first, for each experiment (i.e. KD and breast cell lines), each row was multiplied by the ratio of the median of all row means over the individual row mean. This transformation normalized the row means across each experiment to become the overall median of row means. In the second step, column values were multiplied by the ratio of the average column median across all KD and breast cell lines over the individual column medians. This transformation normalized the column medians of all samples across both experiments to become equal to the global column median. After normalization, the KD and breast cell line tables were concatenated and log2 over median transformed (as described above). The normalized data for the 41 breast cancer cell lines was selected as the background dataset and used to generate the baseline co-regulation network. The baseline protein interaction map was generated by calculating all protein-protein pairwise spearman correlations and the top 16747 positively correlating protein pairs with a Benjamini_Hochberg corrected p-value <0.001 were used to create the co-regulation map.

To get an accurate measure of dysregulations, each knockdown experiment sample dataset was individually combined with the 41 breast cell line data, and bivariate outlier tests for deviations of protein concentration co-regulations was determined for the 16747 protein pairs as described previously (*11*). In brief, dysregulated interactions were identified as bivariate outliers (samples which deviated from the best fit diagonal line) for each protein-protein association expression map. The Mahalanobis distance (MD) (*39*), a multidimensional Z-score, was calculated for each data point. In order to identify outlier samples, the two-tailed Grubbs Outlier test (*40*) was applied in an iterative manner to the MD scores and generated a p-value for every sample supporting an interaction. These p-values were corrected using the Benjmaini-Hochberg method (*41*) and any sample with an adjusted p-value < 0.1 was defined as significantly dysregulated. However, significant outliers may be identified along the best fit protein-protein trend line due to the nature of Mahalanobis distances, therefore a secondary filter was applied which removed dysregulated interactions of samples that appeared within the (1) top 12.5% of expression values for both interacting proteins or (2) bottom 12.5% of expression values for both interacting proteins. These false positive dysregulations were high (or low) expressors for both interaction members.

The UBR4 knockdown induced deviations were determined for the two shRNA probes (1 and 2) as well as a control (CTL) for each of the 8 CNI high and 7 CNI low cell lines. Using a sequence of nested filters we isolated the sets KD induced deviations specific to each cell line. Beginning with the unfiltered MD values for each cell line, we calculated the coefficient of variation (CV) for each interaction between the KD sample datasets. Interactions with a CV <= 25% between KD samples were retained and their KD MD values were averaged. Following the CV filter the log2 fold change between the averaged KD MD and the control MD for each interaction was calculated. Those interactions with a KD/CTL MD ratio of at least 2-fold were isolated and any interaction that passed the two previous filters in more than one cell line were removed. Finally, the remaining interactions were filtered based on their Grubbs Outlier test adjusted p-value (FDR<0.25). For each high |CNI| cell line UBR4 depletion-induced protein expression changes of interacting proteins (log2 protein_1 / protein_2) were compared to their underlying CNA (CNA protein_1 - CNA protein_2) values for all significantly dysregulated interactions, to determine convergence or divergence of protein and CNA levels upon UBR4 depletion.

All data analyses and statistical tests were performed using R (*42*).

### Index of Supplementary Tables

**Table S1.** Gene copy number alteration (CNA), RNA, and protein levels of 36 breast cancer cell lines, correlations of CNA, RNA, and protein levels, and correlations of protein and RNA levels of multi-protein complex members (Fig. 1).

**Table S2.** Protein-protein interactions used for mean copy number imbalance (|CNI|) determination and |CNI| levels for 668 cell lines (*13*) (Fig. 2a and fig. S2).

**Table S3.** Correlation of gene dependency and |CNI| (Fig. 2b and fig. 2)

**Table S4**. *In vitro* and *in vivo* cancer cell line viability data upon UBR4 depletion (Figs. 2d-f, fig. 3)

**Table S5.** Proteome maps of 16 cancer cell lines upon UBR4 depletion and control treatment.

**Table S6.** Gene set enrichment analysis of proteins differentially regulated in high and low |CNI| cancer cell lines upon UBR4 depletion (Fig. 3)

**Table S7.** Deviations of protein co-regulations upon UBR4 depletion (Fig. 4)

**Table S8.** Protein concentration changes and CNA levels in protein interactions dysregulated in high |CNI| cell lines upon UBR4 depletion (Fig. 4)

**Figure S1.**
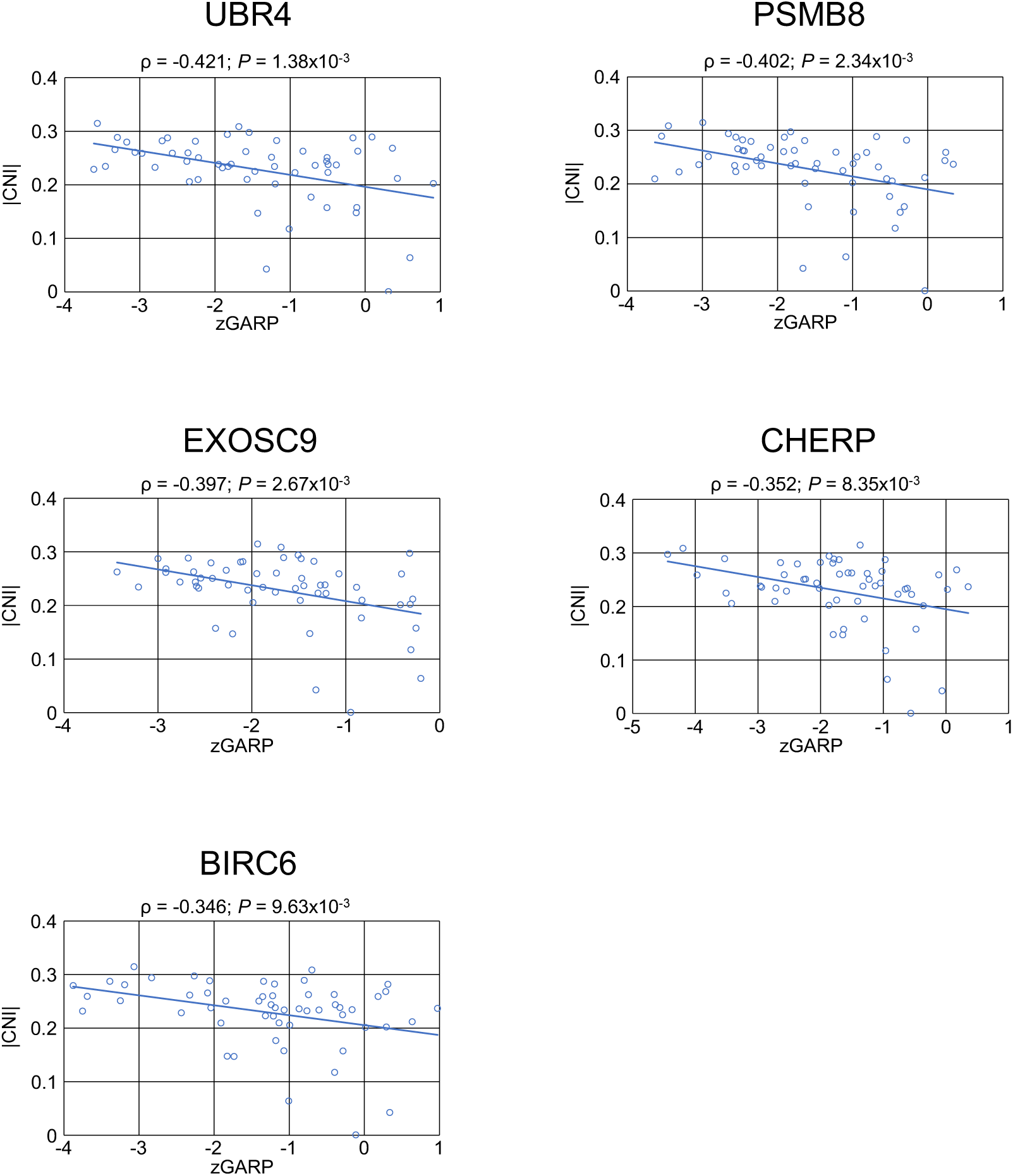
Correlations of |CNI| and gene essentiality (determined using RNAi technology) for UBR4, PSMB8, EXOSC9, CHERP, and BIRC6, across 55 breast cancer cell lines (*16*) (table S2) (ρ, Pearson’s correlation coefficient; *P*, p-value determined using the R Pearson cor.test function (*42*), zGARP and Dependency Score are measurements for gene dependencies from Marcotte et al. (*16*).

**Figure S2.**
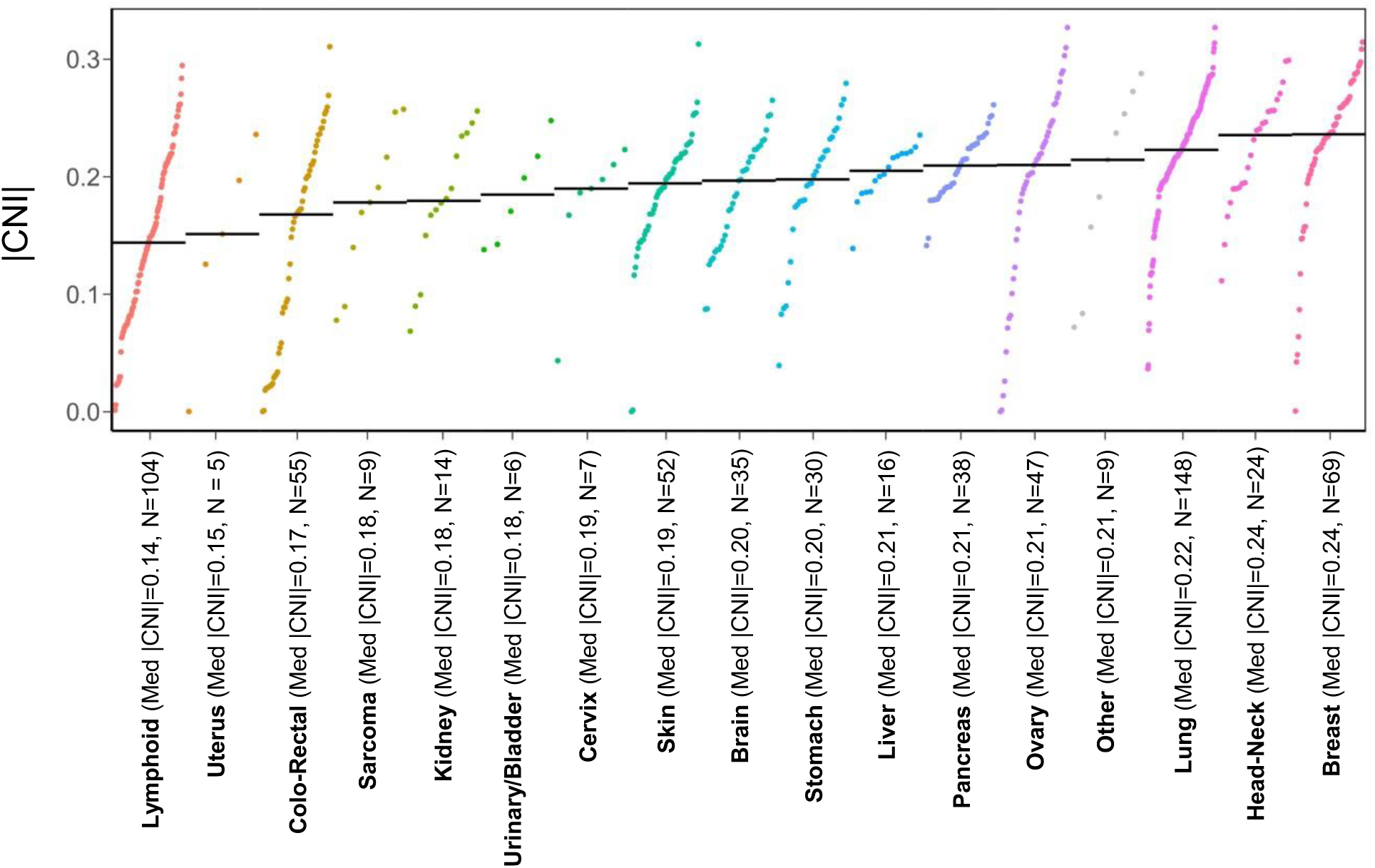
|CNI| levels for 668 cell lines (*13*) (table S2; the category “Other” comprises adrenal, pleura, prostate, and testis as tissues of origin; median |CNI| values for all cell lines for each tissue of origin and number of samples in the categories are indicated in parentheses).

**Figure S3.**
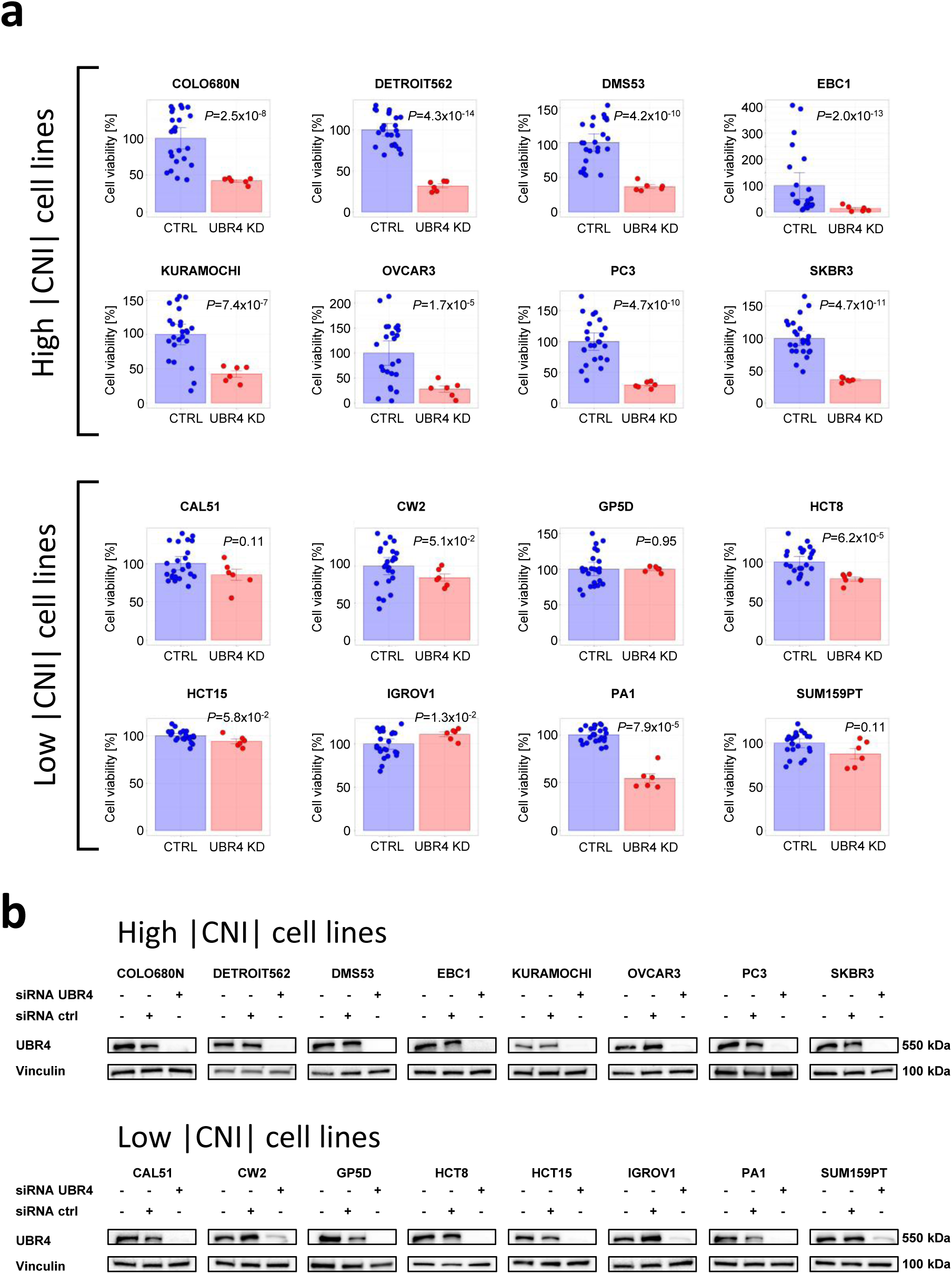
*In vitro* cell viability data upon UBR4 depletion in 16 cancer cell lines. (**a**) Cell lines were treated with a pool of UBR4 targeting siRNA (red) or unspecific control siRNA (four individual probes, blue) for 6 days and cell viability was determined by measuring intracellular ATP concentration (six replicates, significance was evaluated using the Student’s t-test, two-tailed, unequal variance). Average ratios are shown in Fig. 2d. High |CNI| cell lines (COLO680N, DETROIT562, DMS53, EBC1, KURAMOCHI, OVCAR3, PC3, SKBR3) are shown in the upper two rows, and low |CNI| cell lines (CAL51, CW2, GP5D, HCT8, HCT15, IGROV1, PA1, SUM159PT) in the lower two rows. (**b**) Western blot data showing UBR4 depletion in all 16 cell lines for untreated controls and upon 6-day treatment with a pool of UBR4-targeting or unspecific siRNA probes.

**Figure S4.**
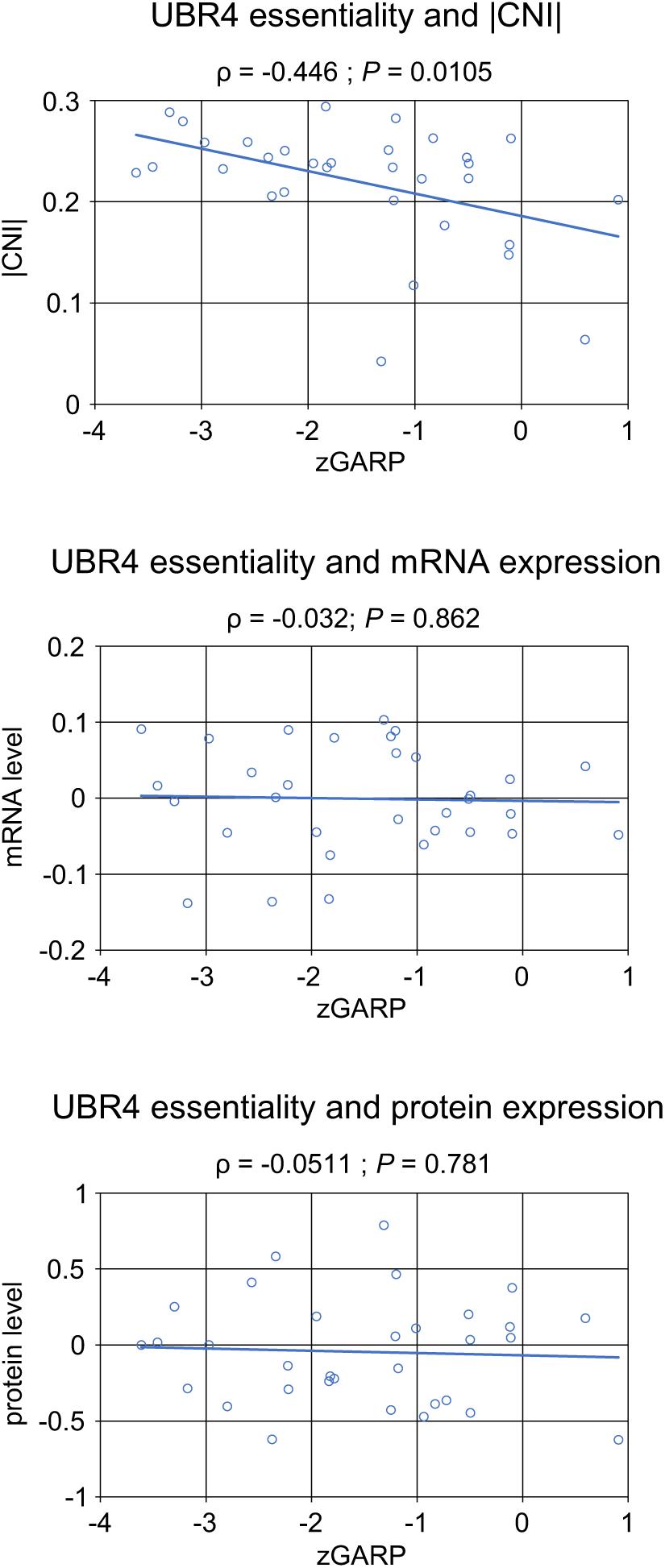
UBR4 dependency in breast cancer cell lines is significantly correlated with |CNI| (top plot, also see fig. S1), but not with mRNA expression (middle) or protein expression (bottom). The plots show data across 32 breast cancer cell lines with available gene dependency (*16*), mRNA (*13*), and proteome data (*11*). UBR4 dependencies were derived from a genome-wide short hairpin (shRNA) loss-of-function screen and are reported in normalized Gene Activity Ranking Profile (zGARP) units (*16*), protein and mRNA levels are shown as log2 ratios over the median of all 32 cell lines.

**Figure S5.**
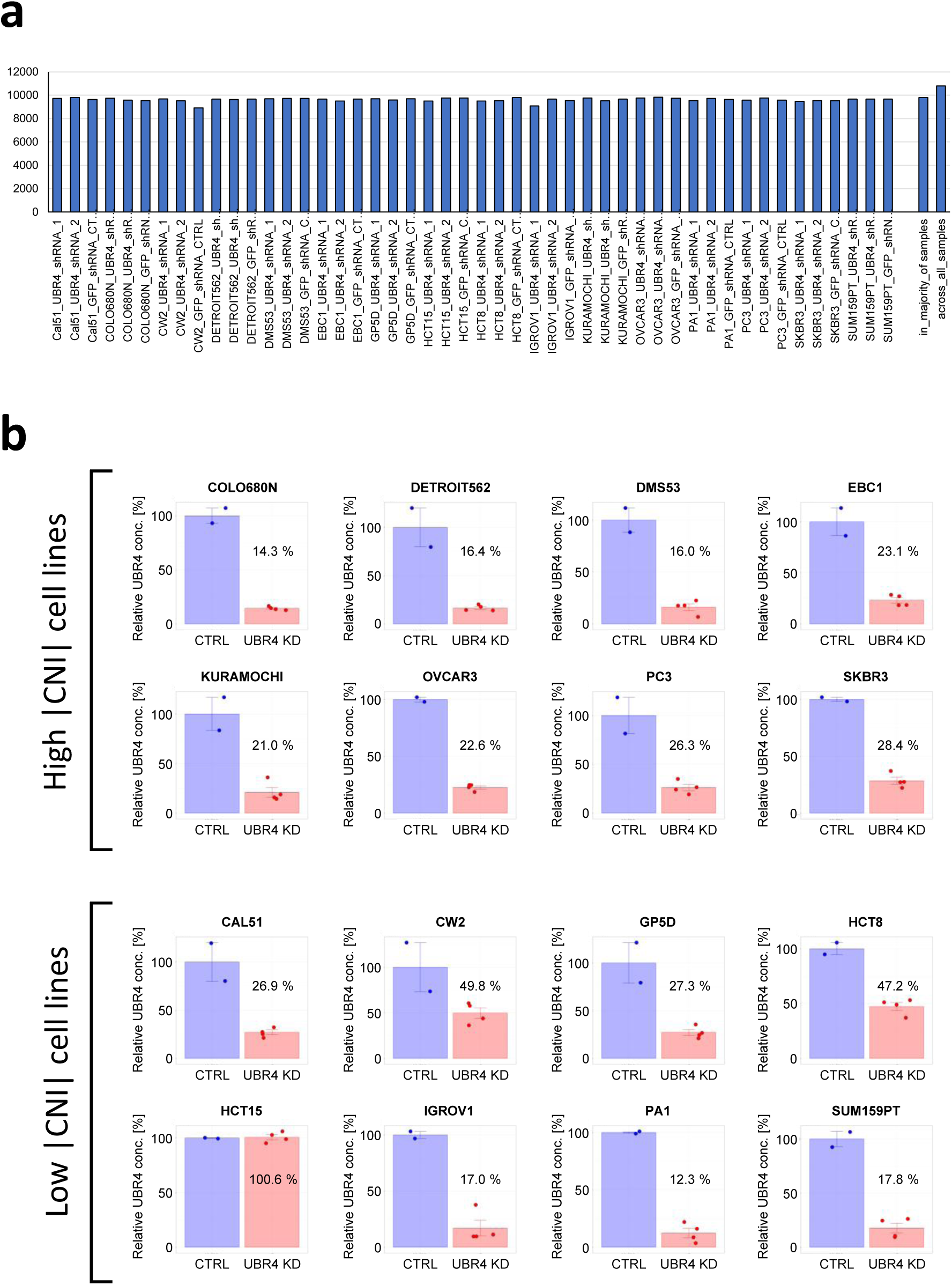
Proteome maps of 16 cancer cell lines upon shRNA-based UBR4 depletion and control treatment. (**a**) Numbers of proteins quantified in each sample using multiplexed quantitative mass spectrometry-based proteomics (UBR4 depletion was achieved with two different shRNA probes (1 and 2), control samples were generated through stable expression of GFP-targeting shRNA; two replicates were analyzed for each sample and replicate data were averaged (see Methods); on average 9625 proteins were quantified per sample, 10796 proteins were quantified across all samples, and 9804 in the majority of the samples). (**b**) UBR4 levels in each sample upon UBR4 depletion and control treatment were determined by multiplexed quantitative proteomics. Average relative UBR4 levels compared to controls are indicated in each plot. No UBR4 depletion was measured for HCT15 and these data were excluded from further analysis.

**Figure S6.**
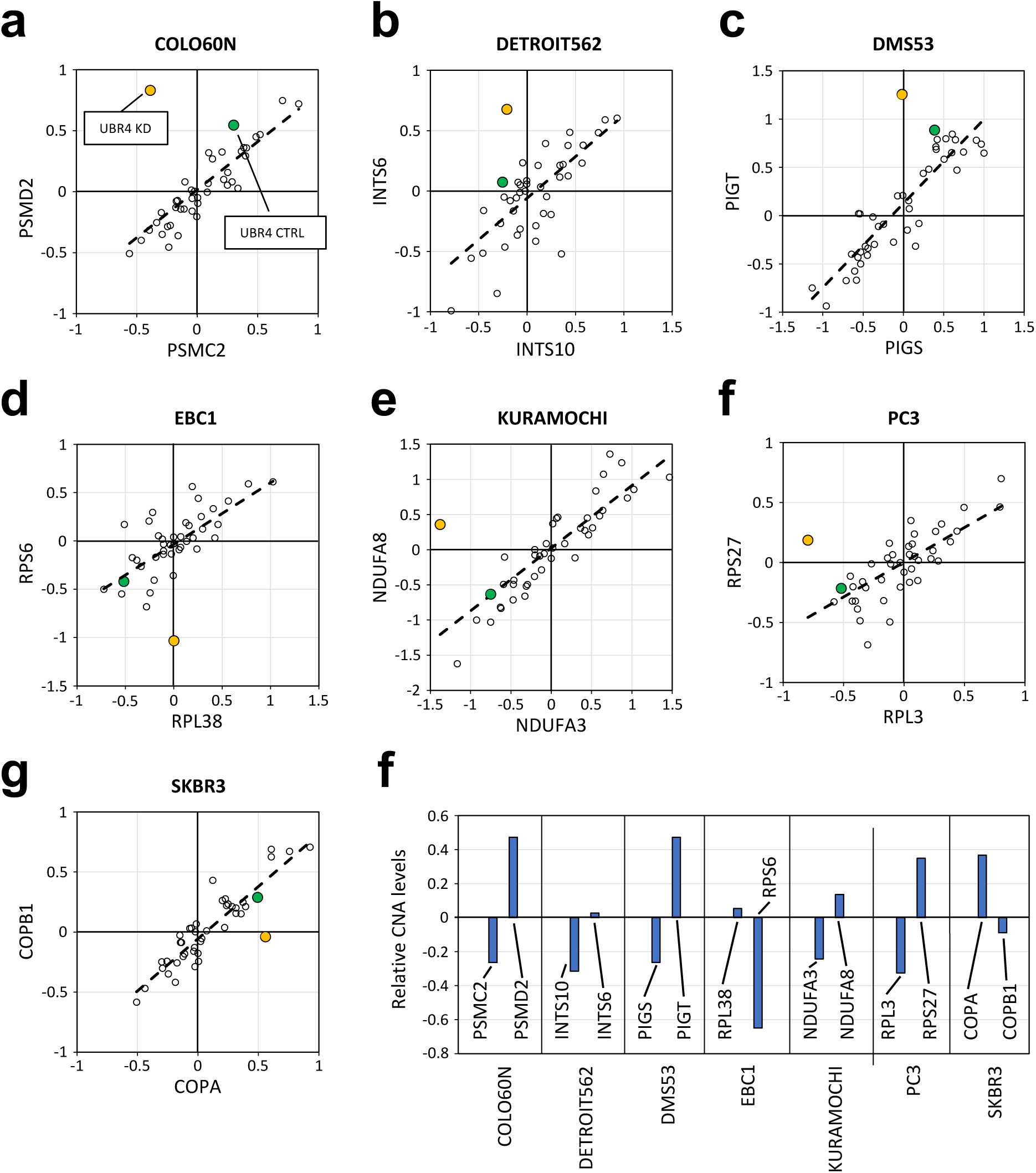
(**a**-**g**) Examples of UBR4 depletion-induced protein interaction dysregulations in high |CNI| cell lines (see Fig. 4b for an example from the cell line OVCAR3). Cell line names are on the top of the plots, protein concentrations are shown as log2 ratios over the median concentration of all 16 cell lines analyzed in this study and 41 breast cancer cell lines (*11*). The protein concentration in the high |CNI| cell lines upon UBR4 depletion are shown in orange, those in the control samples in green. The other points plotted represent levels of the proteins at baseline across the 41 breast cancer cell lines. (**f**) Relative CNA levels (log2 over median of all genes) for the protein pairs and cell lines shown in (**a-g**) (*13*).

